# Spatial Transcriptomic Analysis Reveals HDAC Inhibition Modulates Microglial Dynamics to Protect Against Ischemic Stroke in Mice

**DOI:** 10.1101/2024.08.08.607139

**Authors:** Kevin Jayaraj, Ritesh Kumar, Sukanya Shyamasundar, Thiruma V Arumugam, Jai S Polepalli, S Thameem Dheen

## Abstract

Ischemic stroke significantly contributes to global morbidity and disability through a cascade of neurological responses. Microglia, the immune modulators within the brain, exhibit dual roles in exacerbating and ameliorating ischemic injury through neuroinflammatory and neuroprotective roles, respectively. Despite emerging insights into microglia’s role in neuronal support, the potential of epigenetic intervention to modulate microglial activity remains largely unexplored. We have previously shown that sodium butyrate, a histone deacetylase inhibitor (HDACi) epigenetically regulates inflammatory response of microglia after ischemic stroke and this study was aimed to characterize the transcriptomic profiles of microglia and their spatial distribution in the stroke brain followed by HDACi administration. We hypothesized that the administration of HDACi epigenetically modulates microglial activation and a region-specific microglial phenotype in the stroke brain, shifting their phenotype from neurotoxic to neuroprotective and facilitating neuronal repair and recovery in the ischemic penumbra. Utilizing a rodent model of middle cerebral artery occlusion (MCAo), spatial transcriptomics and 3D morphometric reconstruction techniques were employed to investigate microglial responses in critical penumbral regions, such as the hippocampus, thalamus, cortex and striatum following HDACi administration. We found that HDACi significantly altered the microglial transcriptomic landscape involving biological pathways of neuroinflammation, neuroprotection and phagocytosis as well as morphological phenotype, promoting a shift towards reparative, neurotrophic profiles within the ischemic penumbra. These changes were also associated with enhanced neuronal survival and reduced neuroinflammation in specific regions in the ischemic brain. By elucidating the mechanisms through which HDAC inhibition influences microglial function, our findings propose therapeutic avenues for neuroprotection and rehabilitation in ischemic stroke, and possibly other neurodegenerative conditions that involve microglia-mediated neuroinflammation.

## Introduction

Ischemic stroke is one of the primary causes of long-term disability and mortality worldwide, presenting a significant public health challenge by exacerbating both health and economic burdens on society (Donkor, 2018; Feigin *et al*., 2022). Focal ischemic stroke which is characterized by abrupt cessation of blood supply to a specific region in the brain, leads to hypoxia (a deprivation of oxygen), triggering a cascade of deleterious cellular and molecular events such as excitotoxicity, neuroinflammation, and apoptosis. During reperfusion, restoration of blood flow to the infarcted tissue further aggravates tissue injury (Traystman, 2003; Mergenthaler, Dirnagl and Kunz, 2022). Complex, multicellular pathways and interactions dictate mechanisms that mediate neuronal damage post-ischemic injury and highlight the need for novel therapeutic strategies for enhancing recovery and clinical outcomes (Zhang *et al*., 2019; Wang, Leak and Cao, 2022).

Microglia, resident immune cells of the brain are at the forefront of the body’s response to ischemic injury. They exhibit a spectrum of phenotypic changes (Dubbelaar *et al*., 2018; Wendimu and Hooks, 2022) and transition dynamically between conventional pro-inflammatory (M1) and anti-inflammatory (M2) states depending on spatial and temporal contexts after ischemia. In the stroke brain, microglia initially clear debris and release pro-inflammatory cytokines like TNF-α and interleukins, triggering neuroinflammatory responses crucial for initiating repair but turn deleterious if it progresses without biological checkpoints controlling the response (Wang, Leak and Cao, 2022). Transitioning to an anti-inflammatory phenotype, microglia release cytokines and neurotrophins promoting tissue repair and limiting ongoing damage (Gupta *et al*., 2018; Zhang *et al*., 2021; Charlton *et al*., 2023). The balance between these microglial states and their heterogenous expression profiles are crucial for the prognosis of stroke. Positive modulation of microglial activation is essential in ameliorating neurodegeneration and reperfusion mediated injury, suggesting that targeted interventions aimed at microglia could offer significant therapeutic benefits (Nagy *et al*., 2020).

Epigenetic mechanisms, such as DNA methylation, histone modification, and RNA-associated silencing, play pivotal roles in regulating microglial function and response to ischemic stroke. Histone deacetylases (HDACs) are key enzymes involved in removing acetyl groups from histones, leading to reduced gene expression that has been explored to regulate progress of pathology and neuroinflammation in stroke and other inflammatory conditions (Adcock, 2007; Fessler *et al*., 2013; Patnala *et al*., 2017). In the context of ischemic stroke, such epigenetic modifications influence the phenotypic switching of microglia between pro-inflammatory and anti-inflammatory states, thus affecting the overall inflammatory milieu and recovery process (Saw *et al*., 2020; Qiu, Xu and Zhan, 2021).

Sodium butyrate, a HDAC inhibitor (HDACi), has emerged as a promising epigenetic therapy for ischemic stroke (Sun *et al*., 2015). By modulating the epigenetic landscape, HDACi can shift microglial activation towards neuroprotective states, reducing neuroinflammation and promoting neuroprotection. This therapeutic potential is further supported by studies demonstrating its ability to enhance the expression of neurotrophic factors and modulate key signaling pathways involved in cell survival, neurogenesis, and synaptic plasticity (Patnala *et al*., 2017; Ziemka-Nalecz *et al*., 2017; Saw *et al*., 2020; Li *et al*., 2024; Xu *et al*., 2024).

Our research leverages advanced methodologies, including high-resolution imaging, 3D microglial reconstruction, and spatial transcriptomics, to delve deeper into the effects of HDACi on microglial dynamics and their transcriptional profile in the stroke brain. By exploring biological networks relevant to microglia-neuron interaction and the impact of epigenetic modulation on microglial function, we aim to uncover novel therapeutic targets that could significantly enhance the functional outcomes in ischemic stroke therapy (Qiu, Xu and Zhan, 2021; Zhang *et al*., 2022)

Given the intricate interplay between microglia, neurons, and other glial cells in the ischemic brain, understanding the precise molecular pathways including those affected by epigenetic modifications induced by microglial activity, is critical for developing targeted therapies for stroke. The application of HDAC inhibitors like sodium butyrate offers a unique approach to modulating the brain’s innate immune response, steering it towards pathways that support neuroprotection, repair, and regeneration. Such insights are crucial for developing targeted interventions which harness the therapeutic potential of microglial modulation in the context of ischemic stroke.

The present study demonstrates that HDACi treatment not only reduces infarct volume in stroke brain but also influences microglial phenotypes towards reparative states, facilitating neuronal survival and mitigating neuroinflammation. Through analysis incorporating cutting-edge spatial transcriptomic techniques, we obtain nuanced understanding of microglial roles in stroke pathophysiology and recovery, paving the way for targeted therapies that can capitalize on the therapeutic promise of microglial modulation.

## Methods

### Ethics Statement

Adult male C57Bl/6J mice, 12-15 weeks old were acquired (InVivos Pte Ltd, Singapore) and housed under controlled temperature, humidity, light and dark cycles at the NUS Comparative Medicine (CM) Vivarium. Mice were provided access to standard food pellets and water ad libitum. Animal work was performed in accordance with the Institutional Animal Care and Use Committee (IACUC) through protocol NUS/IACUC/R19-0596 approved for the experiments performed with animal(s) and tissues in this study.

### Surgical Ischemic Stroke Modelling and HDACi treatment in Mice

MCAo modelling was performed as previously described (Patnala *et al*., 2017) using adult male C57BL/6J mice (12-15 weeks old, 26-29g). The mice were anesthetized with isofluorane in an induction chamber. Ophthalmic lubricant (Duratears 3.5g, Alcon, USA) was applied to prevent corneal drying, and body temperature was monitored and maintained at 37.0 ± 0.5°C with a heating pad. The neck of the mice was shaved, disinfected with 70% ethanol and povidone-iodine solution, and a midline incision was made to expose the common carotid artery (CCA), internal carotid artery (ICA), and external carotid artery (ECA). The CCA and ICA were clamped, and the ECA was cauterized and retracted. A Doccol 6-0 monofilament nylon suture (catalog no.: 602123PK10, Doccol Corporation, USA), (diameter of 0.21-0.23 mm), was introduced through the ECA and advanced into the ICA to occlude the MCA. Occlusion was maintained for 60 minutes, after which the suture was withdrawn to allow reperfusion. The ECA incision was cauterized, and the microvascular clips were removed to restore blood flow through the CCA and ICA. The surgical site was sutured, and the animals were allowed to recover from anaesthesia under observation. In sham group mice, surgical reflection of muscles to reveal CCA bifurcation was performed without disturbing the vasculature to affect cerebral blood flow. Sodium butyrate (HDACi) was administered intraperitoneally (I.P.) to the mice (Cat. no. B5887-1G, Sigma-Aldrich) at a dose of 300 mg/kg bodyweight. Two doses of NaBu were administered, 1-hour prior ischemia and 6-hours post reperfusionas described previously (Patnala *et al*., 2017). Mice were housed a temperature-controlled environment with a 12-hour light/dark cycle and provided food and water ad libitum. Motor deficit after surgical modelling was evaluated through preliminary neurological deficit scoring (pNDS). Severity of behavioural deficit and rescue was plotted for each functional parameter of circling, latency to move home cage length to enrichment shell, spasticity/paralysis and allodynia (n=3 in Sham and MCAo and n=4 in MCAo+HDACi groups) (Garcia *et al*., 1995; Li *et al*., 2000). Animals were regularly monitored for adverse effects or complications, ensuring compliance with guidelines for the care and use of laboratory animals, aimed at minimizing suffering and the number of animals used. 24 hours post-stroke induction, mice were euthanized with an intraperitoneal overdose of sodium pentobarbital (150 mg/kg) and absence of pedal and corneal reflexes was confirmed prior to harvest of brain tissue. For histological and immunofluorescence analyses, mice were perfused with ice-cold 4% paraformaldehyde (PFA) and post-fixed at 4°C for 24 hours prior to histological processing. For spatial transcriptomic analyses, brain tissue was processed as per 10x Genomics Visium fresh frozen tissue preparation guide CG000240 - Rev E.

### TTC Staining of mouse brain tissue and evaluation of tissue damage

For 2,3,5-Triphenyltetrazolium chloride (TTC) staining, fresh brain tissue was sectioned into 2 mm coronal slices using a brain matrix. The brain slices were submerged in cold PBS to maintain tissue viability. Freshly prepared 2% solution of TTC (Cat. no. T8877, Sigma-Aldrich) was added to brain sections in a 24-well plate and incubated at 37°C for 30 minutes in a dark, humidified incubator. Viable tissue stained red, while infarcted areas remained pale or white. Post-incubation, sections were rinsed with PBS and transferred to a clear, flat transparent sheet for digital imaging on a flatbed scanner (CanoScan LiDe 400, Canon). Image processing and analysis of TTC-stained brain sections was performed by thresholding scanned images to greyscale and extrapolating pixel values (Goldlust *et al*., 1996).Scanned images were loaded and processed using R packages to calculate and distinguish viable (non-infarcted) and non-viable (infarcted) area in brain tissue. Viable and non-viable areas were calculated in square millimeters based on pixel counts and known image resolution. Statistical analysis, including t-tests and ANOVA, was performed to compare the groups (n=3).. The results were then imported into Microsoft Excel for visualization.

### Histological Processing of Mouse Brain Tissue Sections

Harvested brain tissues were post-fixed in 4% paraformaldehyde and dehydrated through a graded series of ethanol baths and clearene. Dehydrated and cleared samples were then incubated in molten paraffin wax, embedded and allowed to solidify at room temperature. Solidified wax blocks were mounted onto a microtome, trimmed, and sectioned at 7μm thickness. Sections were floated in a warm water bath, transferred onto microscope slides, and stained with toluidine blue. Routine staining protocols were performed involving deparaffinization in clearene, rehydration through descending ethanol series, staining in toluidine blue followed by rinsing, dehydrating ethanol series and tissue clearing prior to coverslipping slides with mounting medium (S302380-2, Dako North America Inc.). For cryosectioning, PFA-fixed brain tissues were cryoprotected in 30% sucrose until fully saturated, embedded in OCT compound, and rapidly frozen using dry ice or pre-cooled isopentane. Frozen OCT-embedded tissue blocks were sectioned at −20°C to −25°C in a cryostat (Leica Instruments GmbH, Germany), with sections cut at 30 to 50 microns, collected onto chilled slides, and stored at −80°C in sealed containers with desiccants until further use.

### Immunofluorescence and Morphometric analysis

Mouse brain tissues were fixed with 4% paraformaldehyde, cryoprotected in 30% sucrose and embedded in OCT compound. Mouse brains were sectioned at 30 or 50 micron thickness on a cryostat (Leica Instruments GmbH, Germany). The sections were permeabilized with 0.3% Triton X-100 and non-specific binding sites were blocked by incubating sections in 5% normal donkey serum (D9663-10ML, Sigma-Aldrich) or 1% bovine serum albumin (126615-25MLCN, Merck). The sections were then incubated with primary antibodies (antibody list in table below) diluted in blocking solution, overnight at 4°C. After rinsing with PBS, sections were incubated with fluorescence-conjugated secondary antibodies for 1 hour at room temperature, protected from light. Following this, sections were rinsed with PBS and mounted with anti-fade fluorescence mounting medium (ab104135, Abcam). The stained sections were imaged using a confocal microscope (FluoView 1000 and FluoView 3000, Olympus Corporation, Japan) at 10x, 20x, 60x and 100x objectives. Z-stacks were captured at 60x and 100x objectives with a z-plane of 20-30µm with z-sections of 0.5µm. Microglia-labelled confocal, Z-stacked imaging datasets were imported into IMARIS v9.5 software for 3D reconstruction and analysis of microglial morphology (n=3). The ‘Surpass’ view was used to inspect the z-stack, and a 3D volume rendering of the z-stack was created using the ‘Volume’ button. For morphometric evaluation and analysis, the ‘Surfaces’ function was employed by selecting the appropriate marker channel (IBA1 for microglia), setting the ‘Surface Detail Level’ and ‘Background Subtraction’, and using the ‘Thresholding’ slider to separate cells from the background. The surface creation was finalized to generate surfaces encompassing the 3D structures of the labelled cells. Morphometric analysis was exported in the ‘Statistics’ tab, quantifying parameters such as volume and surface area. Images and videos of the 3D reconstructions were exported using the snapshot and animation features in IMARIS v9.5. Comparisons of these parameters between groups (Sham, MCAo, and HDACi-treated) were conducted to evaluate the effects of ischemic stroke and HDACi treatment on microglial morphology.

### Primary Antibodies

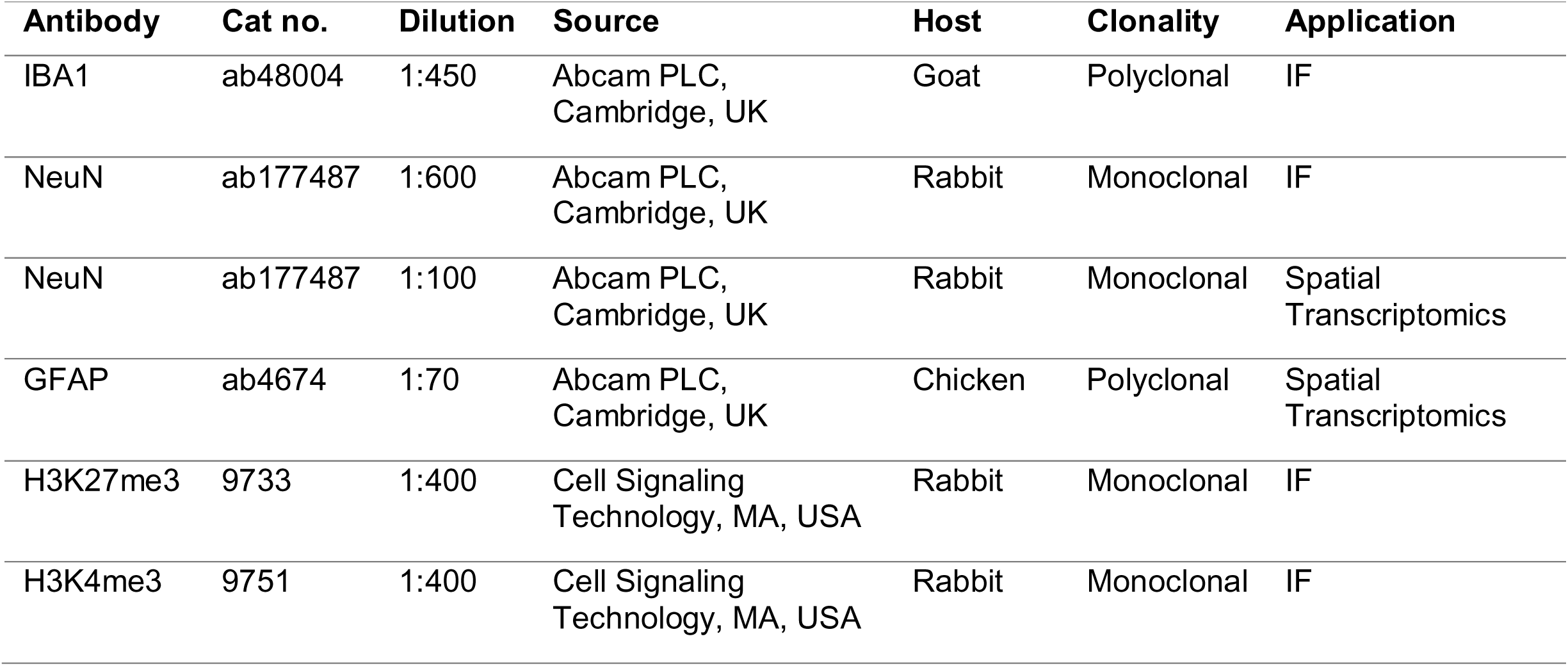

### Secondary Antibodies

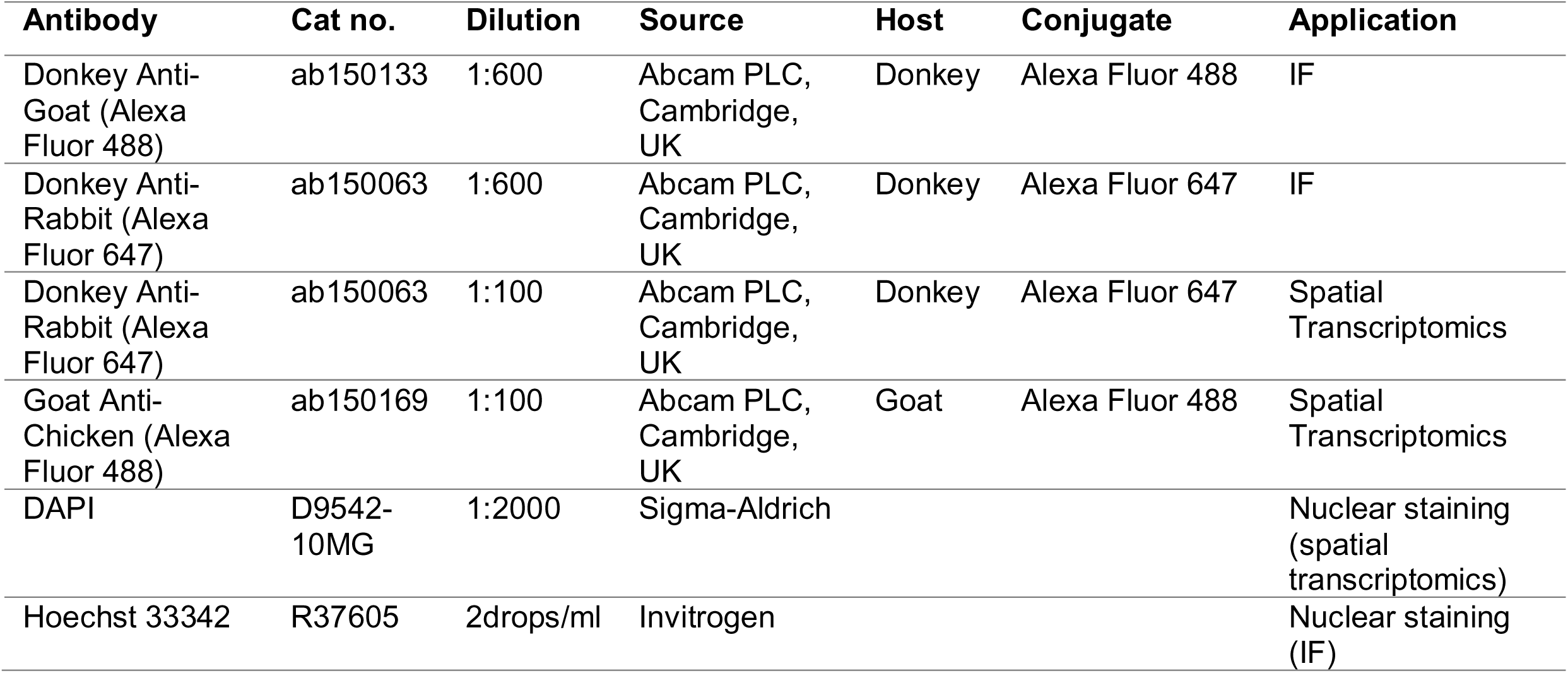

| Antibody | Cat no. | Dilution | Source | Host | Conjugate | Application |
| --- | --- | --- | --- | --- | --- | --- |
| Donkey Anti-Goat (Alexa Fluor 488) | ab150133 | 1:600 | Abcam PLC, Cambridge, UK | Donkey | Alexa Fluor 488 | IF |
| Donkey Anti-Rabbit (Alexa Fluor 647) | ab150063 | 1:600 | Abcam PLC, Cambridge, UK | Donkey | Alexa Fluor 647 | IF |
| Donkey Anti-Rabbit (Alexa Fluor 647) | ab150063 | 1:100 | Abcam PLC, Cambridge, UK | Donkey | Alexa Fluor 647 | Spatial Transcriptomics |
| Goat Anti-Chicken (Alexa Fluor 488) | ab150169 | 1:100 | Abcam PLC, Cambridge, UK | Goat | Alexa Fluor 488 | Spatial Transcriptomics |
| DAPI | D9542-10MG | 1:2000 | Sigma-Aldrich |  |  | Nuclear staining (spatial transcriptomics) |
| Hoechst 33342 | R37605 | 2drops/ml | Invitrogen |  |  | Nuclear staining (IF) |

### Spatial Transcriptomics Pipeline

Fresh-frozen brain tissue samples from the three groups (n=3) were sectioned at 10 μm thickness using a cryostat (Leica Instruments GmbH, Germany). Tissue optimization was performed with TO slides (part no. 3000394, 10x Genomics, California, USA) and reagents as per 10x Genomics user guide CG000238-RevE and involved testing permeabilization durations, with six minutes selected based on tissue morphology preservation and RNA quality, assessed with a Bioanalyzer (Bioanalyzer 2100 system, Agilent, Santa Clara, CA, USA) (Figure 4 Supplementary Figure 1, 2). The sections were placed on Visium Spatial Gene Expression slides (part no. 2000233, 10x Genomics, California, USA) ensuring precise alignment and adhesion to capture areas for spatial mapping. Immunofluorescence staining (as per 10x Genomics user guide CG000312-RevD) was performed using antibodies against neuronal nuclei (NeuN) and glial fibrillary acidic protein (GFAP), with DAPI for nuclear staining, to correlate spatial gene expression data with specific cell types and anatomical features. Antibody concentrations and incubation times were optimized through preliminary titration experiments as advised by 10x standard protocols, to ensure specificity and maximal signal-to-noise ratios. Following immunofluorescence and tissue preparation, in situ reverse transcription on the Visium Spatial slides was performed to convert localized mRNA into cDNA while retaining spatial coordinates using unique molecular identifiers (UMIs) and spatial barcodes. Library preparation, conducted according to Visium protocols (10x Genomics user guide CG000239-RevF), included second-strand synthesis, adaptor ligation, and amplification steps. Libraries were quantified and validated for quality using Novogene’s (NovogeneAIT Genomics, Singapore Pte Ltd) sequencing pipeline for their Illumina platform, to ensure suitability for high-throughput sequencing. Sequencing was performed to a depth of 50 million reads to cover the complexity of the brain transcriptome and ensure spatial resolution of gene expression patterns. Sequenced spatial libraries were processed using 10X Genomics SpaceRanger 2.0.1, aligning reads to the reference genome and attributing them to specific spatial locations on tissue sections based on their spatial barcodes. Spatial transcriptomic data was exported from Loupe Browser 6.0 into Microsoft Excel for consolidation, upload and visualization; systematically organizing metrics such as Log2 Fold Change (Log2FC) values for genes of interest across different tissue sections and experimental conditions. Data wrappers such as venny (Oliveros, 2007), S.R. Plot (Tang *et al*., 2023) and heatmapper.ca (Babicki *et al*., 2016) were utilized for downstream data processing and visualization. The Kyoto Encyclopedia of Genes and Genomes (KEGG), the Database for Annotation, Visualization, and Integrated Discovery (DAVID), and Gene Ontology (GO) profiler were employed to enrich and categorize gene expression data within biological pathways, functional annotations, and cellular component. Data was visualized through S.R Plot (Tang *et al*., 2023) with clusterprofiler and pathview packages in the form of enrichment bubble and ontology bar plots. Venny, an interactive tool for comparing lists (Oliveros, 2007) was utilized to generate comparative lists of upregulated, downregulated and rescued genes across the biological groups with spatial segregation of microglial transcriptomes. Heatmaps were generated using Heatmapper.ca (Babicki *et al*., 2016) to visualize Log2FC data exported from MS Excel organized into selected pathways from enrichment analyses, providing a graphical representation of gene expression variations across different brain regions and treatment groups.

### Statistical analyses

For infarct area evaluation from TTC stained brain slices, one-way ANOVA (F-statistic=32625.11, p=2.74 × 10^-9^) and independent two-sample t-tests (<1×10^−6^) were performed on the calculated non-viable (infarcted) areas from all three biological groups (n=3). The high F-statistic value confirms that the variability between group means is much larger than the variability within groups while the t-tests confirm the findings from one-way ANOVA. Similarly, independent two-sample t-tests revealed significant differences (<0.05) in IMARIS morphometry data focusing on microglial surface area; microglial volume exhibited significant differences (<0.005) as depicted in data between Sham and MCAo, as well as between MCAo and MCAo+HDACi groups. Spatial transcriptomics data processed using proprietary 10x software, SpaceRanger 2.0.1 and Loupe Browser 6.0 utilizes a statistical framework based on sSeq for differential expression analysis (Yu, Huber and Vitek, 2013), Moran’s I for spatial enrichment (Getis, 2007) and Benjamini-Hochberg correction for controlling false discovery rates (p<0.05) natively in the transcriptomics pipeline as described in 10x Genomics repositories. Online data wrappers were fed downstream data exported from Loupe Browser 6.0 by filtering for genes limited to the selected pathways within the microglial/neuronal spatiotranscriptomes by MS Excel’s VLOOKUP function and data was visualized in the form of heatmaps (heatmapper.ca), enrichment bubble plots, combined Gene Ontology bar plots, CNETplots for KEGG pathways-targets enriched (S.R. Plot) and Venn diagrams for dysregulated gene expression across microglial spatiotranscriptomes (Venny).

## Results

### HDACi treatment reduces ischemic infarct volume in the brain and enhances preliminary locomotor function of mice post-ischemic insult

HDACi was administered intra-peritoneally to mice undergoing 1-hour ischemia and 24-hour reperfusion periods in two doses. The first dose at 1-hour pre-ischemic insult to prime cells and the subsequent dose post 6-hours of ischemic insult to effectively influence reperfusion mediated injury. As reported earlier, MCAo was performed for 1h and reperfusion post-ischemia was done for 24h prior to euthanasia (Fig.1A) (Patnala et al., 2017). Neurological viability and infarct volume were assessed through toluidine blue (Nissl) staining and TTC (2,3,5-Triphenyltetrazolium Chloride) (Fig.1B-C). From the staining, the extent of ischemic penumbra at brain regions of interest (ROI)’s such as hippocampus, striatum, thalamus and the cortex were identified to proceed with assessment of microglia-neuron interaction in regions (Fig.1B). Evaluation of basic locomotor activity in mice prior to euthanasia (Fig.1D, Fig.1 Supplementary Video 1) showed circling contralateral to the infarct, and reduced mobility in the MCAo groups, while HDACi-treated mice did not exhibit circling and showed reduced latency to move within the home cage towards enrichment shell. Substantive restoration of viable neuronal soma was observed (Fig.1E-F) 24-hour after administration HDACi. Taken together, the findings suggest a) successful modelling of ischemia in the mouse brain; along with identification of regions of interest (ROIs) (hippocampus, thalamus, striatum and cortex) in the ischemic penumbra, presenting as potential areas to target reperfusion mediated neuronal damage. b) Gross evaluation of viable brain tissue and preliminary behavioural performance of HDACi administration after MCAo showcases a potential restorative effect, substantiating a need for further assessment of microglial contribution to ischemia-reperfusion mediated neuroinflammation as well as neuroprotection.

**Figure 1:**
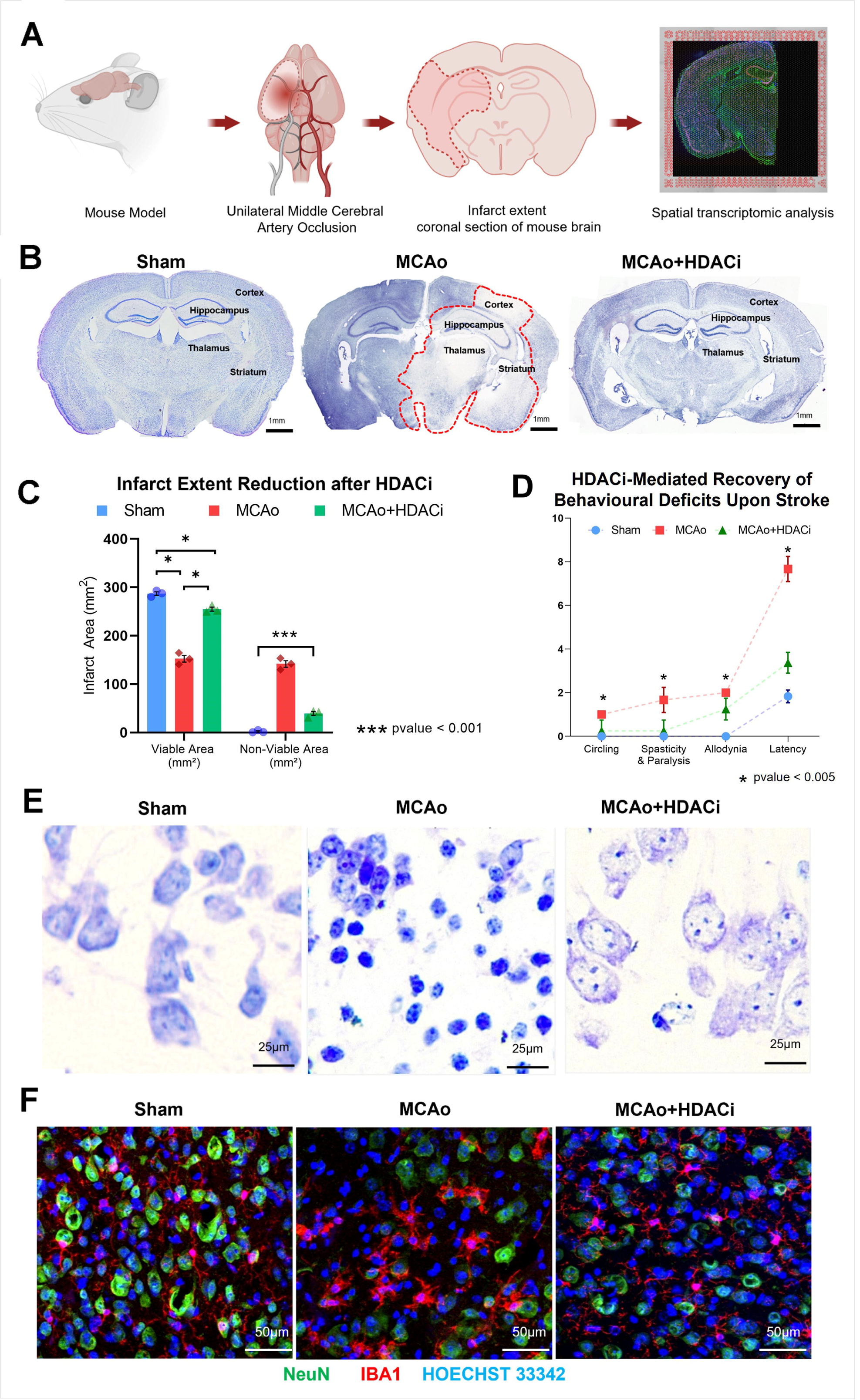
Middle cerebral artery occlusion and gross neurological status upon HDACi treatment. (A) Graphical pipeline describing ischemic stroke modelling in the rodent brain, ischemia-associated microglia and phenotypes. (B) Toluidine blue (nissl) staining of brain tissue sections with MCAo and Sham showcasing infarct extent and anatomical regions of interest (ROI) covered under the penumbra. (C) Epigenetically mediated protection through HDACi attenuates infarct volume 24hours post ischemic insult (TTC or 2,3,5-Triphenyltetrazolium Chloride assessment of brain slices exhibit substantial recovery of viable brain tissue upon NaBu administration) (n=3 in each experimental group). (D) Behavioural assessment of ischemic affect through neurological deficits occurring through stroke demonstrates enhanced locomotor activity upon HDACi administration with reduced circling, spastic paralysis, allodynia and locomotor latency (to reach enrichment shell within home cage) after HDACi treatment (n=3 in Sham and MCAo and n=4 in MCAo+HDACi groups). X axis represents behavioural paradigm tested. Y axis represents scoring achieved for each group - Circling: [no = 0, yes = 1], Latency: [1 to 8 seconds to reach enrichment shell] Spasticity and Paralysis: [No spasticity or paralysis = 0, Spasticity = 1, Spastic paralysis = 2] Allodynia: [no allodynia or vocalization = 0, allodynia sensitivity = 1, allodynia+vocalization = 2]. (E) Toluidine blue stained mouse brain sections of dentate gyrus neurons in the Hippocampus exhibit variance in infarct intensity and neuronal damage post-ischemic stroke with HDACi administration. (F) Immunostained confocal images of NeuN (channel AF 674, false-colored green) and IBA1 (channel AF488, false-colored red) appear to display preserved neuronal organization in the penumbral cortex and reduced microglial aggregation at the infarct periphery upon HDAC inhibition. Nucleus is stained with Hoechst 33342 (blue).

### HDACi ameliorates neurodegeneration in brain nuclei after ischemic stroke and inhibits microglial aggregation

The hippocampal CA3-dentate gyrus are particularly vulnerable regions of ischemic insult (Einenkel and Salameh, 2024). Histological evaluation of toluidine blue stained sections of the hippocampal CA3-dentate gyrus for neuronal and Nissl body aggregation, reveals a degree of neuroprotection after HDACi administration (Fig.1E); preservation of Nissl bodies correlates with enhanced neuronal survival (Smith et al., 2012), substantiating that HDACi confers neuroprotection in vulnerable regions of the mouse brain. Confocal images (Fig.1F) showing immunofluorescence expression of NeuN and IBA1 in neurons and microglia respectively, in the cortex surrounding infarct periphery indicate recruitment of microglia in stroke brain while there was a reduction in recruitment and alteration of microglial morphology in stroke brain following HDACi treatment; implying a tempered inflammatory response that mitigates further neuronal damage by fostering a more conducive environment for recovery (Lee et al., 2014).

### HDACi treatment inhibits ischemia-induced microglial phenotype in mouse brain

Microglial cells that are ramified with slender processes in the sham mouse brain exhibit amoeboid and reactive phenotypes in the stroke brain (Fig.2A). Upon HDACi administration, microglial phenotypes shift towards a sham spectrum ramified phenotype with long and slender processes (Fig.2A). Quantitative morphometric analysis by 3D-reconstruction of thick tissue sections generated using IMARIS image analysis software (v9.5) reveals that microglial surface area decreases while intracellular volume is increased in the stroke brain compared to sham brain microglia (Fig.2B-C). However, HDACi treatment reverses the stroke microglial phenotype closer to sham mice, suggesting that HDACi inhibits ischemia-induced phenotypic changes in microglia (Fig.2B-C). The distinct microglial morphology observed in this study represents transcriptomic profiles that may influence their role in inflammatory response and neuroprotection through the course of pathophysiology after ischemic stroke.

**Figure 2.**
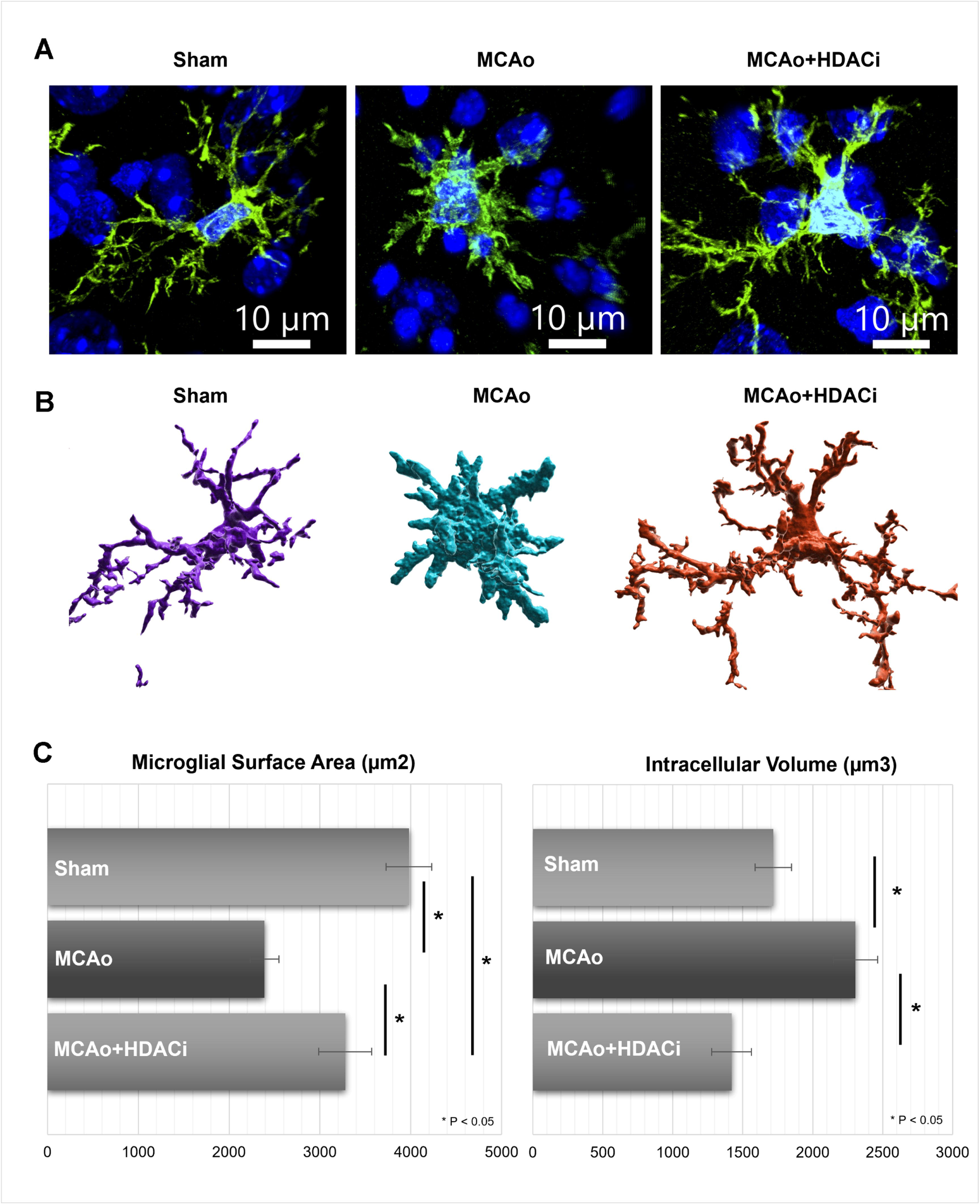
Microglial morphometric assessment through surface reconstruction with IMARIS 9.5. (A) High-resolution confocal images of Microglia stained with IBA1 (channel AF 488, green) reveal diverse morphological phenotypes between biological groups (n=3 in each experimental group). (B) Individual microglial cells were segmented and processed across groups to obtain surface area and cellular volume morphometric data. 3D-reconstruction of microglial surfaces spotlights microglia with ramified and long, slender processes in HDACi inhibitor administered mouse brain reminiscent of sham brain microglia while MCAo brain microglia are amoeboid with substantial depth across the z-stack. (C) Microglial surface area and intracellular volume are inverse between stroke and epigenetically treated groups, presenting a phenotypic reversal of microglia in the penumbra.

### HDACi inhibits ischemia-induced increase in nuclear expression of histone marks H3K27me3 and H3K4me3 in microglia

Phenotypic heterogeneity of microglia in pathological conditions is regulated by epigenetics, and various signaling molecules and transcription factors (Wang *et al*., 2021). We have previously shown that acetylation marks such as H3K9ac regulate the inflammatory response of microglia in the stroke brain (Patnala *et al*., 2017). In the present study, we have analyzed the expression pattern of methylation marks such as H3K27me3 and H3K4me3 to understand molecular mechanisms contributing to microglial phenotypic heterogeneity in the ischemic brain. The nuclear expression of H3K27me3 (Fig.3A-B) and H3K4me3 (Fig.3C) appeared to be remarkably elevated in microglia of the stroke brain. Upon HDACi administration, microglia exhibit a more ramified phenotype with inhibition in nuclear expression of H3K27me3 and H3K4me3, similar to expression patterns of sham brain microglia suggesting that microglial phenotypic heterogeneity is influenced by histone mark status. Interestingly, we also found neuronal cells in the hippocampal CA2-CA3 gyri (Fig.3A-B) exhibiting an inverse H3k27me3 expression pattern to microglia, suggesting an epigenetic feedback mechanism across microglia-neuron regulatory network of histone status that warrants further study.

**Figure 3.**
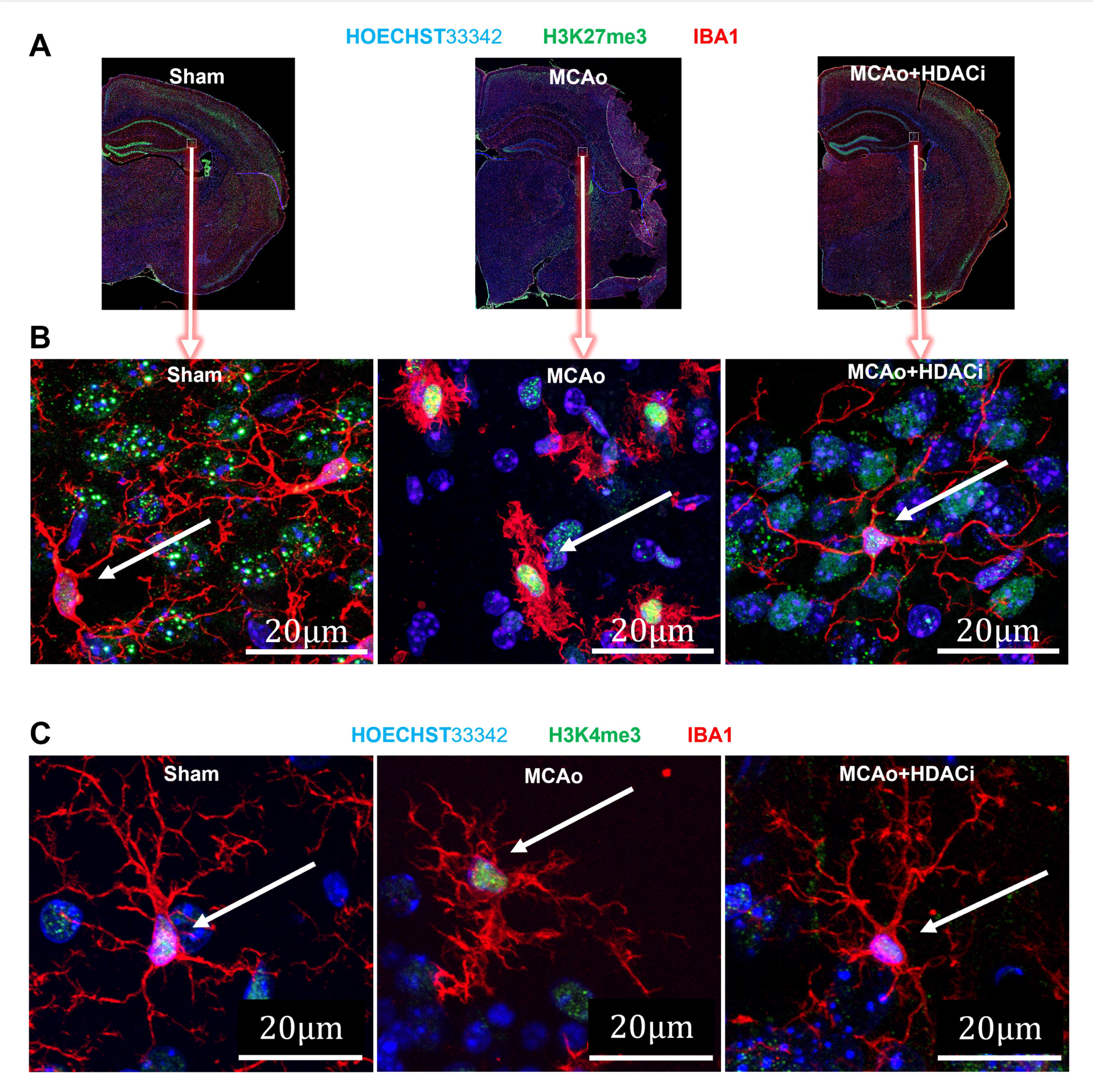
HDAC inhibition and histone mark status in microglia after ischemic stroke. (A-B) Hyper-intense expression of H3K27me3 (channel 647, false-colored green) in MCAo brain microglia (IBA1, channel 488, false-colored red) appear to be rescued towards sham expression levels in hippocampal microglia from HDACi-treated stroke brains. Potential shuttling of H3K27me3 suggests upstream involvement of the PRC2 complex and EZH2, components mediating H3K27me3-related gene silencing upon ischemic insult. (C) Similar to H3K27me3, H3K4me3 (channel 647, false-colored green) expression levels in sham and HDACi-treated stroke brain tissue showcase similar expression patterns with stroke elevating expression in microglial nuclei suggesting neuron-microglia regulatory networks activated by HDACi administration.

### Mapping of spatial transcriptome of microglia in ischemic mouse brain following HDACi administration

The spatial transcriptome profile of microglia in response to ischemic injury and HDACi administration was analyzed using the 10x Genomics Visium (FF) platform. Fresh frozen brain slices were immunolabelled and imaged for neurons (NeuN, red) and astroglia (GFAP, green) along with DAPI (blue) on GEX Visium slides prior to library preparation for the spatial overlay (Fig.4A-B). Immunoflourescence imaging and proprietary Loupe Browser v6.0 cluster overlay with key ROIs, namely the hippocampus, thalamus, striatum, and cortex were marked for post processing (Fig.4A). Stroke brain tissue reveals a loss of cellular heterogeneity, and distinct organizational patterns present in sham brain tissue, suggesting the disruption of cellular microenvironments and signaling networks upon ischemic insult (Fig.4B-C). Preliminarily, the spatial transcriptomic data revealed 19 distinct cell populations based on marker gene expression across the three experimental groups (Fig.4F) with a Uniform Manifold Approximation and Projection (UMAP) plot demonstrating the spatial distribution of these populations alongside the spatial overlay (Fig.4E, 4C). To specifically isolate the microglial transcriptome, a sub-clustering approach through feature selection and thresholding in loupe browser v6.5.0 was employed (Fig.4D) by leveraging a list of microglia signature genes identified in recent studies (Gerrits *et al*., 2020; Quintana *et al*., 2022). This allowed for the extraction of microglia-enriched spots from brain co-ordinates to generate distinct microglial spatiotranscriptomic datasets within the four penumbra ROIs.

**Figure 4.**
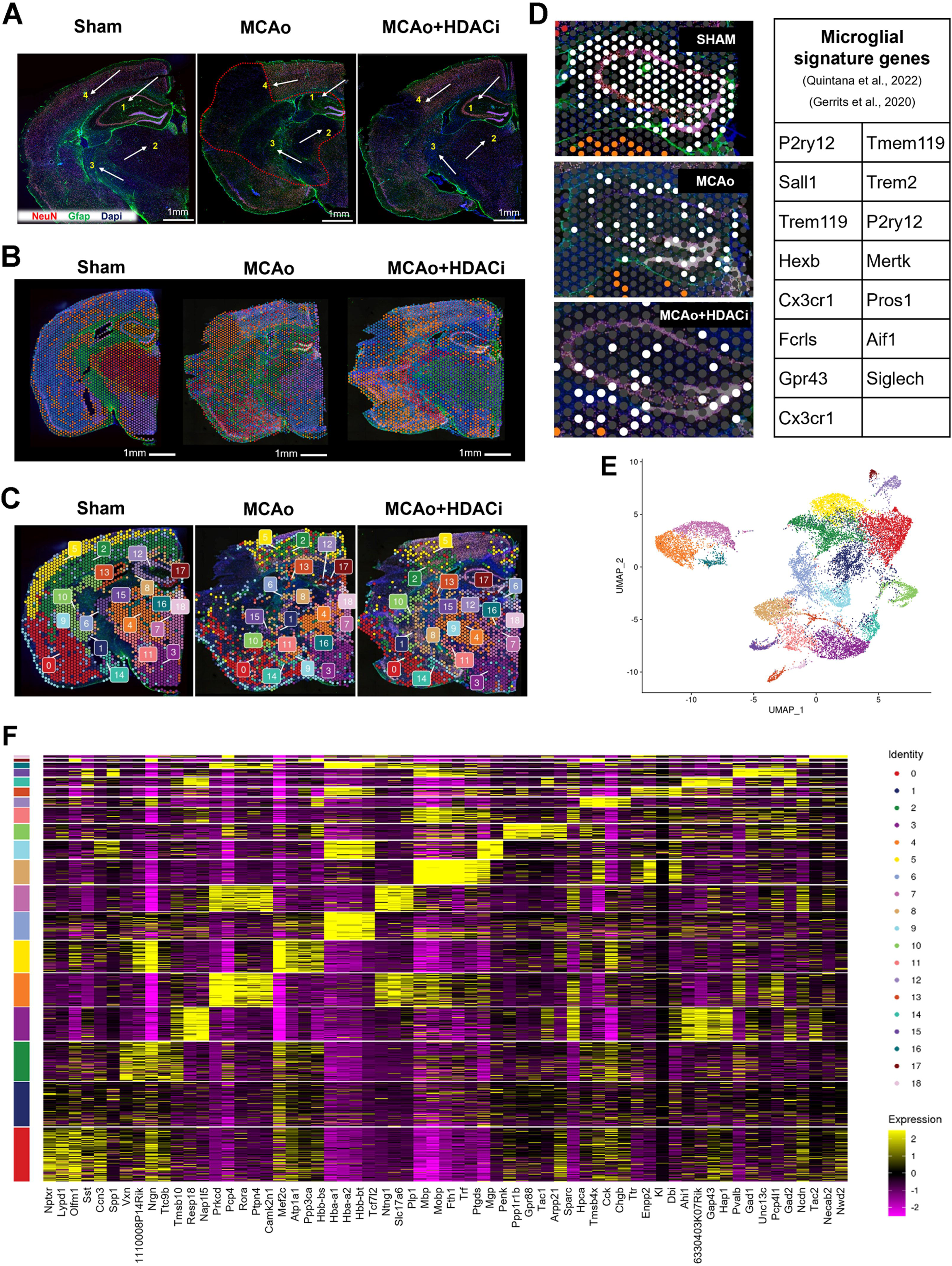
Spatial transcriptome pipeline: ROIs, population clusters and microglial spatiotranscriptomic signature (Visium Fresh Frozen) (A) Immunofluorescence (IF) image and spatial (spot) overlay (B) of Visium fresh frozen sections and marker gene population clusters (C) (n=3 in each experimental group). (B) 10x Genomics proprietary Loupe Browser 6.0’s default transcriptome overlay represented as k-clusters and population identities; (C) the distinct population clusters appear to lose organization upon ischemic insult. (D) Microglial spots enriched through microglia signature genes adapted from (Gerrits *et al*., 2020; Quintana *et al*., 2022) performed through Loupe browser 6.0’s feature selection and thresholding to output microglia-specific spatial transcriptomes segmented based on regions on interest in the tissue section. (E) Heatmap of total spatial library with cell signature markers for processed brain samples exhibiting 19 population identities. (F) Uniform Manifold Approximation and Projection (UMAP) dimension reduction of all cell populations in the dataset.

### HDACi alters microglial spatiotranscriptome profiles following ischemic stroke

Differential expression analysis and pathway enrichment studies were carried out to investigate microglial spatiotranscriptomics and their functional implications. The anatomic regions of interest on the overlay of spatiotranscriptomic spots, focusing on the ischemic penumbra has been demarcated using loupe browser v6.0 (Fig.5A). Four-set Venn diagrams were drawn to identify region-specific, unique and commonly upregulated (Fig.5B) /downregulated (Fig.5C) /rescued (Fig.5D) microglial genes across the ischemic penumbra in the hippocampus, thalamus, striatum and cortex. Among the upregulated microglial genes across the four regions of interest, 22% of genes (3544) was found to be commonly upregulated after HDACi administration (Fig.5B). On the other hand, only 1.5% (190) of microglial genes was found to be commonly downregulated across the four regions of interest (Fig.5C). Additionally, the datasets of differentially expressed genes (both up/downregulated) were processed to obtain microglial genes that were modified or rescued (closer to sham expression levels) (Fig.5D) following HDACi treatment.

**Figure 5.**
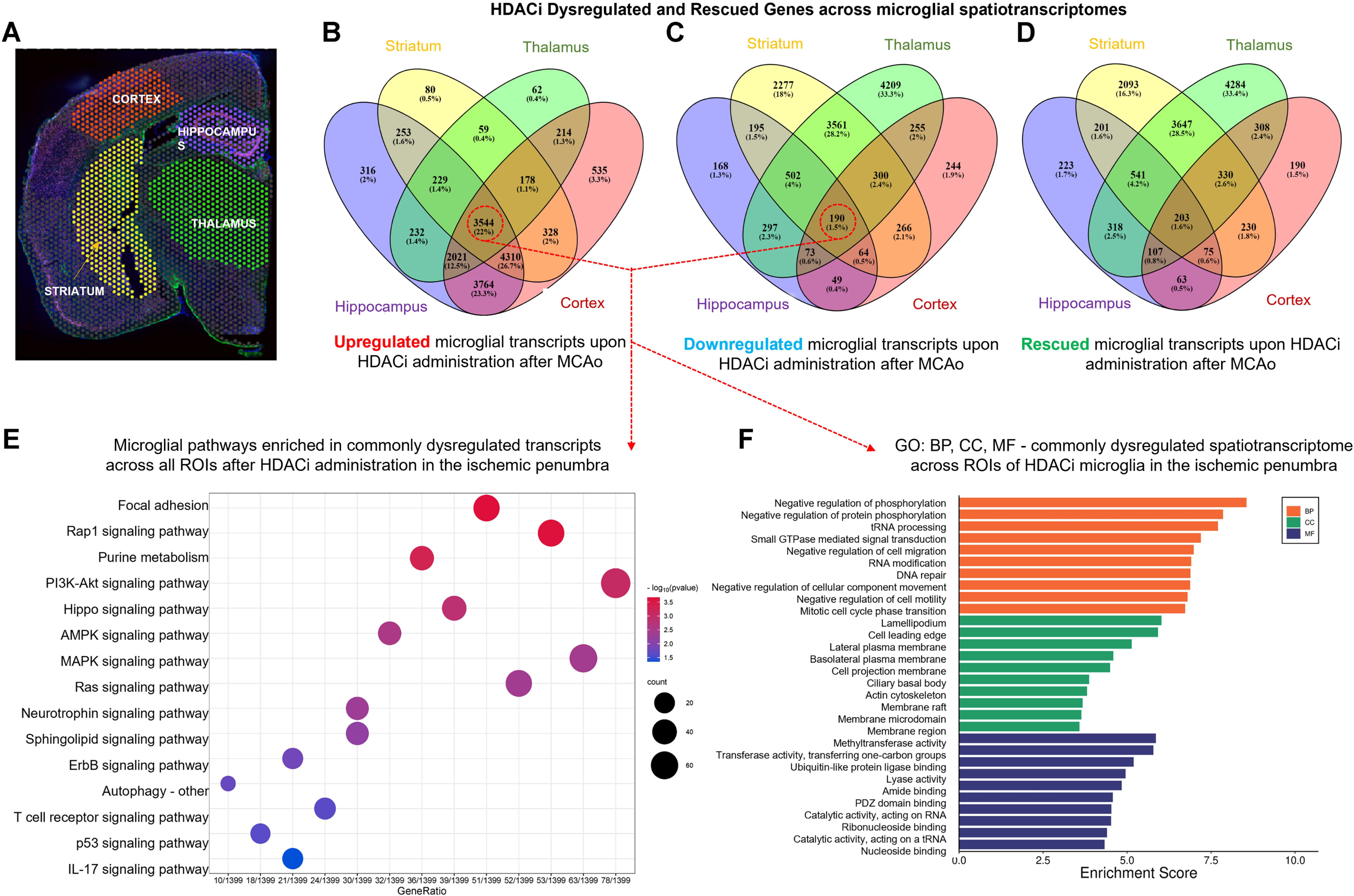
Spatial segmentation of microglial transcriptomes: expression signatures across the ischemic penumbra and enriched pathways (KEGG) (A) Grouping and segmentation of Penumbra regions of interest in *LoupeBrowser* for spatial demarcation of microglia after ischemic stroke. (B, C, D) Four-set Venn diagrams of unique and overlapping HDACi-Upregulated (B), Downregulated (C) and Rescued (D) Genes, elucidate the homogenous and heterogenous marks in microglia across brain regions. Pathway (Kegg) Enrichment (E) and Gene Ontology (F) analysis of genes dysregulated homogenously across the four microglial spatiotranscriptomes are processed and showcase potential microglial pathways of interest to analyze and target after ischemic stroke. (n=3 for each experimental group)

The dysregulated microglial genes were subjected to KEGG pathway enrichment (Fig 5E) and Gene Ontology (GO) (Fig.5F) analyses. Pathway analysis yields enrichment of key functional pathways pivotal for neuroinflammatory response, mediating neurotrophic support, and facilitating phagocytosis (Fig.5E) alongside other pathways that may be relevant to microglial responses after ischemic stroke (Fig.5 Supplementary Figure 1). Regulatory signaling pathways such as PI3K-Akt signaling (PIK3CA, PIK3CB, AKT1, AKT2, AKT3, PTEN), FC gamma R-mediated phagocytosis (FCGR1, FCGR3, FCGR4, SYK, and PIP5K1C), MAPK signaling (DUSP1, DUSP4, DUSP6, ERK1-2, cJUN, UCP4), TNF signaling (TNF, NFKB1, RelA), and TGFβ signaling (TGFBR1, TGFBR2, SMAD2, SMAD3, SMAD4) pathways shed light on the transformative influence of epigenetic regulators on microglial functionality in the post-stroke brain environment. The genes from these pathways were found to be modified closer to sham expression levels in each region of interest within the transcriptome of microglia after HDACi administration.

Thus, HDACi treatment results in a significant alteration of the microglial transcriptomic landscape, with a spatial context. Pathway-enrichment (Fig.6 Supplementary File 1) and GO analyses conducted for the microglial genes of hippocampus, thalamus, striatum, and cortex (Fig.6 Supplementary Figure 1A-1B) revealed that HDACi alters, and rescues expression of several key pathways involved in neuroprotection, inflammatory response and phagocytosis after ischemic stroke (Fig.6A,6B,6C,6D). These analyses of microglial spatiotranscriptomes shed light on the emphasis of these pathways describing their big-picture and region-specific roles after ischemic insult. Heatmaps visualize the expression pattern of rescued transcripts across the functionally relevant pathways, alongside radar plots delineating the top 50 rescued targets from these pathways in each microglial spatiotranscriptome (Fig.6A,6B,6C,6D). Confocal microscopy of microglia in the hippocampus (from Fig.2B), can be corroborated with microglial spatiotranscriptome to draw parallels between HDACi-modified microglial transcriptome <> microglial phenotypic heterogeneity in the hippocampus (Fig 6A <> Fig.2B).

**Figure 6.**
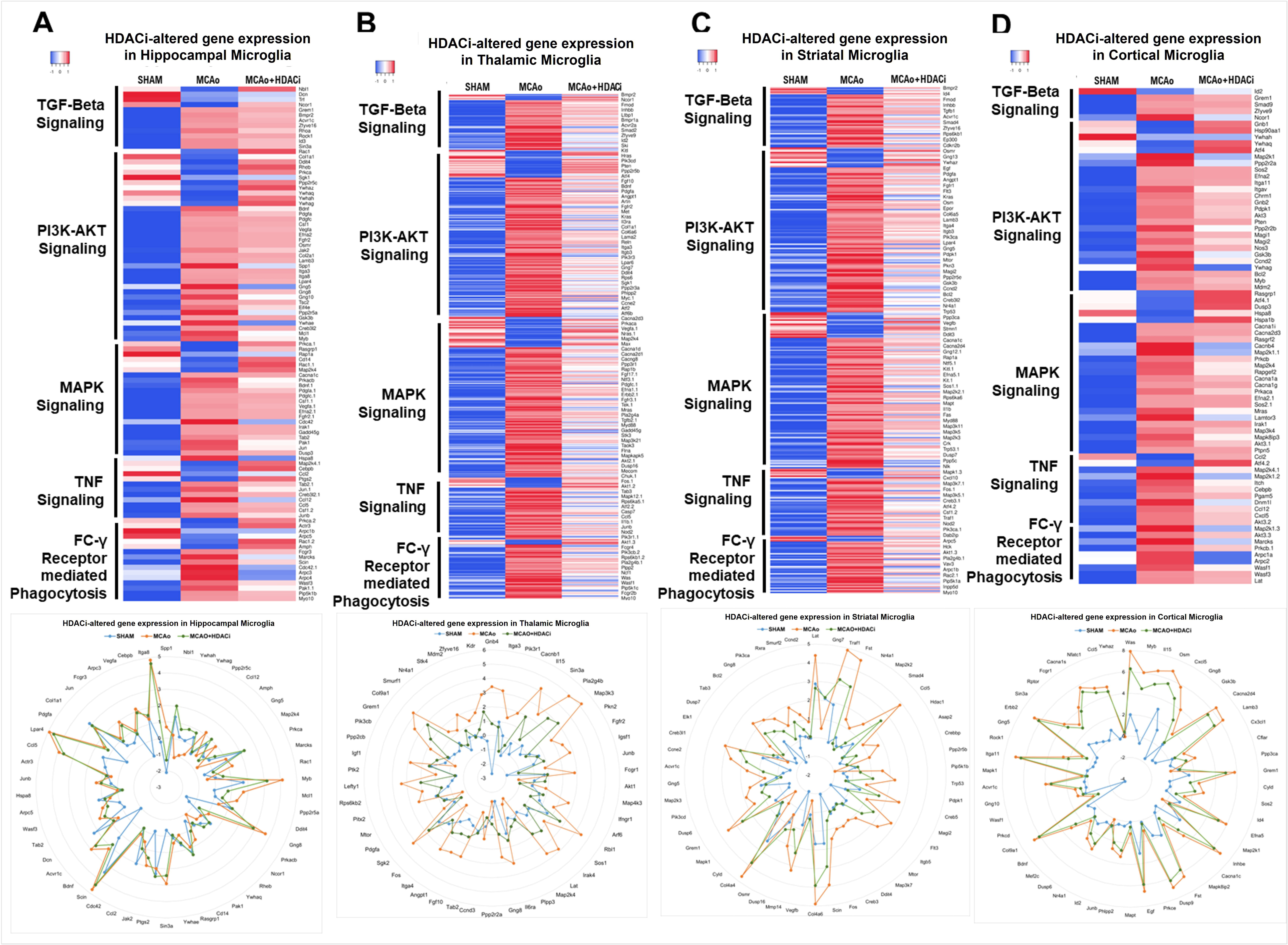
Microglial spatiotranscriptome expression of enriched, key biological pathways after HDACi administration in ischemic stroke. Heatmaps of microglial spatiotranscriptomes of each ROI, hippocampal (A), thalamic (I), striatal (J), cortical (K) microglia exhibit differential profiles of selected biological pathways that were enriched and involved in ischemic injury across the three biological groups (n=3 for each experimental group). Modified and rescued (closer to sham expression) genes within identified pathways crucial for neuroprotective, neuroinflammatory and phagocytosis mediation responses are represented, and top 50 targets are represented in the radar plot for each penumbra ROI.

The spatiotranscriptomic profile of microglia across groups emphasizes the impact of HDACi on microglial gene expression involved in neuroprotection. We also analyzed if HDACi treatment improves neuroprotection by influencing the neuronal spatiotranscriptome in the penumbra following ischemic stroke. The HDACi-modified spatiotranscriptome of penumbral neurons reveal a distinct inverse expression pattern across groups in enriched pathways (Fig.7B,7C), between ischemia-affected and HDACi treated mouse brains (Fig.7A) with broader implications for neuroprotection, synaptic remodeling, and plasticity. This suggests a granular modulatory effect of HDACi that is potentially influenced by the local microenvironment and neuronal signaling to remodel microglial expression profiles.

**Figure 7.**
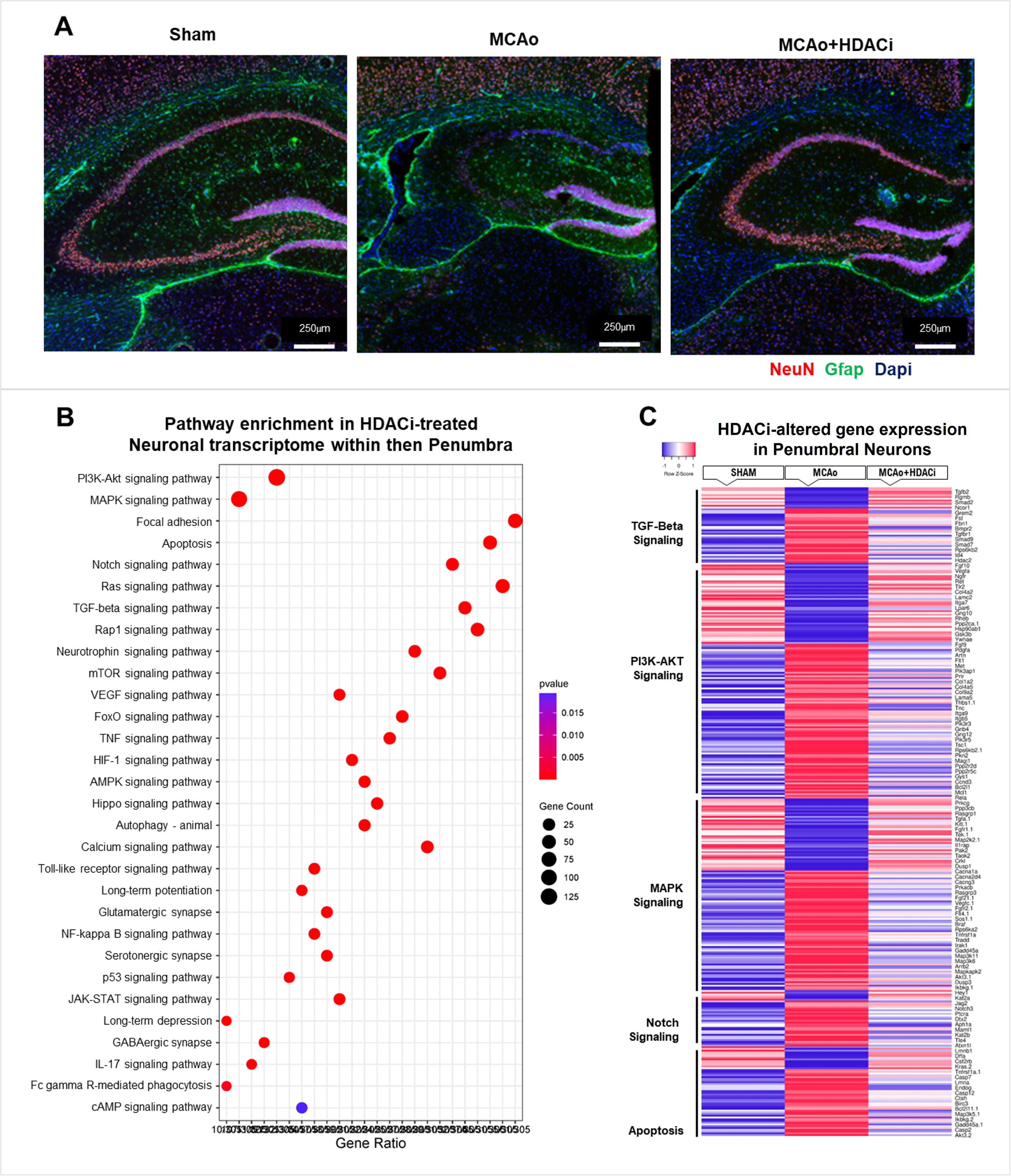
Penumbra Neuronal Spatiotranscriptome. (A-B) Spatiotranscriptome of penumbral neurons showcases enriched neuronal pathways relevant for survival through HDACi-mediated rescue after ischemic stroke. (C) Distinct, inverse expression profiles after HDACi treatment can be observed, indicating context-dependent microglia-neuron interaction upon ischemic stroke and epigenetic regulation through HDACi. (n=3 for each experimental group)

The findings highlight the spatial and molecular heterogeneity of microglial responses to ischemic injury and HDAC inhibition, emphasizing the therapeutic capabilities of epigenetic modulation in ischemic stroke. The identified pathways and rescued genes provide a robust framework for further exploration of microglial-neuron interactions and the development of targeted therapies.

## Discussion

Recent advancements in spatial transcriptomics analysis have enriched our ability to dissect stroke pathophysiology by providing spatial resolution to gene expression patterns in neurons and glial cells including microglia. In the current study, integration of spatial transcriptomics with advanced imaging and reconstruction techniques has offered a comprehensive view of microglial dynamics in the stroke brain. The MCAo model recapitulates human physiology to facilitate investigating microglial responses within the ischemic brain (Chen *et al*., 2009; Ansari *et al*., 2011), in particular the penumbra regions including the hippocampus, thalamus, striatum and cortex. In the MCAo model, HDACi treatment improves sensory-locomotor deficits inflicted on mice post-ischemic stroke as observed in certain behavioural paradigms prior to euthanasia. An extended battery of behavioural tests were not incorporated in the experimental design due to the 24-hour time point that was optimal for capturing a snapshot of reperfusion-mediated neuroinflammation in the stroke brain. As functional improvements upon HDAC inhibition, particularly sodium butyrate has been described previously in rodent MCAo models (Zhou *et al*., 2021), this study was designed to observe the microglial transcriptome specifically during reperfusion in the brain’s ischemic penumbra. Microglial activation in the stroke brain appears to influence the neuronal transcriptome, which is substantiated by the dysregulated transcriptional responses observed in microglia and neurons of the stroke brain. This altered microglial response, which is regulated by various signalling molecules and epigenetic mechanisms, can be ameliorated through epigenetic intervention as observed in the present study.

### The Influence of Transcription Factors on phenotypic heterogeneity of Microglia

Microglial cells are dynamic and exhibit region- and time-dependent phenotypic heterogeneity in response to neuropathological conditions such as neuroinflammation and neurodegeneration including ischemic stroke (Hickman *et al*., 2018; Li *et al*., 2020). In the current study, differential expression of microglial spatiotranscriptomes associated with phenotypic heterogeneity of microglia in response to ischemic stroke, particularly under the modulation of HDAC inhibition was observed. This variation in microglial phenotypes elucidates the interplay of transcription factors, cell-specific receptors and microglia-neuron interaction; outlining the complexity behind microglial polarization such as neuroprotective or neurotoxic phenotypes in the post-stroke brain (Olah *et al*., 2011; Haroon, Miller and Sanacora, 2016; Dubbelaar *et al*., 2018).

In our microglial spatiotranscriptomic analysis, ischemic stroke was found to alter key signalling molecular pathways involved in neurotrophic support, phagocytic activity and neuroinflammation including PI3K-Akt signaling, FC gamma R-mediated phagocytosis, MAPK signaling, TNF signaling, and TGFβ signaling pathways and transcription factors regulating immune response such as NF-κB, STAT1/STAT3, Nrf, and PPAR family (Yang *et al*., 2018; Simpson and Oliver, 2020; Li *et al*., 2024). Epigenetic mechanisms such as post-translational histone modifications provide an additional layer of regulation to elicit cell-specific and stimuli specific immune responses (Allis and Jenuwein, 2016) in the stroke brain.

Post-translational histone modifications determine basic transcriptional rates and the magnitude of gene expression (Bannister and Kouzarides, 2011). Among the two major histone modifications namely histone acetylation and histone methylation, histone acetylation is associated with increased accessibility to transcriptional machinery leading to increased gene expression (Marmorstein and Zhou, 2014), whereas histone methylation is more complex in regulation of gene expression through repression. Different histone modification patterns of specific marks at different sites have been shown to play distinct biological roles; this prompted us to analyze the spatial transcriptome of microglia distributed in infarct penumbra of the stroke brain upon HDACi administration to understand the epigenetic control of microglia-mediated neuropathological outcomes in the stroke brain.

Modification in histone acetylation and deacetylation status, is maintained by histone acetyltransferase (HATs) and histone deacetylase (HDACs) enzymes respectively (Yang and Seto, 2007). Several studies have shown that inhibition of HDAC exerts immunomodulatory effects, specifically suppressing pro-inflammation and inducing anti-inflammatory signaling pathways (Adcock, 2007; Kulthinee *et al*., 2022). We have previously demonstrated the HDACi-specific immunosuppression of pro-inflammatory mediators (TNFα and NOS2) and upregulation of IL10, an anti-inflammatory mediator in microglia in ischemic stroke mice model exerting neuroprotection (Patnala *et al*., 2017).

Dynamic regulation of repressive histone methylation marks, H3K27me3 and H3K4me3 within microglia play a significant role in their response to ischemic injury and subsequent recovery processes (Cheray & Joseph, 2018; Tang et al., 2014). In the present study, markedly elevated expression of these markers were observed in microglia that are distributed in the penumbra of ischemic brain. The increased nuclear expression of these histone marks in microglia appears to reflect a response to the ischemic environment-induced transcriptional activity which may be associated with changes in migration, phagocytosis and phenotypic heterogeneity inferred through the spatial transcriptome of microglia. Remarkably, this ischemia-induced expression of H3K27me3 and H3K4me3 was found to be reverted to near sham levels in microglia upon HDACi treatment, suggesting epigenetic regulation of H3K27me3 and H3K4me3 downstream pathways that influence recovery mechanisms.

Further HDACi-induced suppression of H3K27me3 in microglia of ischemic brain may be associated with activation of its upstream targets such as Enhancer of Zeste Homolog2 (EZH2) (Suppl.tab.1_averaged_MicroglialLog2FC_10X.xslx) a catalytic subunit of the Polycomb Repressive Complex 2 (PRC2) (Ayata *et al*., 2018) which has been shown to play important roles in the epigenetic regulation of phagocytic clearance and pruning of synaptic activity in microglia (Hong, Dissing-Olesen and Stevens, 2016; Ayata *et al*., 2018). Further, since the PRC2 complex, including EZH2 has been shown to regulate TNFα and TGFβ signalling pathways (Penas and Navarro, 2018; Chen *et al*., 2019), the increase in EZH2 expression post HDACi treatment in microglia may be responsible for altered expression of both inflammatory and neurotrophic regulators such as TNFα and TGFβ signalling observed in the current study.

### Epigenetic Regulation over Pathway-Specific Microglial transcriptomes involved in Neuroprotection, Neuroinflammation and Phagocytosis in the Stroke Brain

Impact of HDAC inhibition on microglia in stroke brain suggests spatially defined transcriptomic changes as the results highlight the spatial heterogeneity of microglial transcriptomes within various regions of the stroke brain. Notably this was evident as pathways crucial to neuroprotection in microglia of stroke brain after HDACi treatment exhibit a transcriptomic signature similar to that of the sham brain.

The TGF-β signaling pathway, central to immune modulation and cell differentiation, displayed marked upregulation of TGFβ receptors and SMAD family genes (e.g., TGFβR1, TGFβR2, SMAD 2-4) in microglia of stroke brain following HDACi treatment. In TGFβ signaling, TGFβR-Smad family complexes regulate the expression of TGFβ target genes (Chandra *et al*., 2012; Taylor *et al*., 2017; Du *et al*., 2018) (Lin *et al*., 2017; Zöller *et al*., 2018; Zhu *et al*., 2023); the surge of TGF-β signaling pathway genes suggests an amplified anti-inflammatory and regenerative response in microglia, potentially enhancing their interactions with neurons to support tissue repair and functional recovery in stroke brain following HDACi treatment. HDACi has been found to upregulate many other anti-inflammatory genes including STAT3 and PPARγ indicating an induction of anti-inflammatory response of microglia in the stroke brain (Patnala et al., 2017).

Further, HDACi treatment restored expression of several other TGF-β signaling pathway molecules including BMP and activin receptor membrane-bound inhibitor (*BAMBI*) which interferes with TGFβ or BMP pathway transduction by forming heterodimers with *TGF*β*RI/BMPRI* (Chen *et al*., 2023). The involvement of *BAMBI*, and various BMPs highlights the diverse role of TGF-β signaling through microglia, influencing not only immune responses but also neurogenesis, and synaptic plasticity (Meyers and Kessler, 2017).

It has been well established that microglia contribute to neuroinflammation through excessive secretion of proinflammatory mediators in the stroke brain (Jayaraj *et al*., 2019). The activation of proinflammatory mediators, is regulated by mitogen-activated protein kinases (MAPK) comprising of extracellular signal regulated kinase (ERK), c-Jun N-terminal kinases (JNK), and p38 and nuclear factor kappa beta (NF-κβ) gene expression (Kaminska, Mota and Pizzi, 2016). HDAC inhibition has been shown to modulate the p38 MAPK-MK2 signaling cascade, regulating microglial activation and cytokine production (Kaminska *et al*., 2009; Kaminska, Mota and Pizzi, 2016). In the present study, many of the MAPK signalling pathway genes were found to be altered in microglia following ischemia and HDACi treatment. Further, in microglia of stroke brain the downregulation of c-Jun upon HDACi administration appears to modulate the neuroinflammatory response, promoting neuroprotection (Waetzig *et al*., 2005). This downregulation of cJun could be mediated by HDACi-induced activation of MKP-1 (DUSP1) which has been shown to inactivate MAP Kinases (Eljaschewitsch *et al*., 2006). Overall, HDACi appears to shift microglial activation towards neuroprotective states, reducing neuroinflammation and promoting neuroprotection through epigenetic modifications.

Further, HDACi-induced upregulation of PI3K-AKT pathway in microglia of stroke brain appears to associated with neuroprotection since we have previously demonstrated that microglial PI3K-AKT-BDNF pathway regulates memory and learning in neurons via long term potentiation (LTP) (Saw *et al*., 2020). In addition, HDACi treatment seems to potentiate an array of neuroprotective signaling cascades such as through the upregulation of microglial TrkB, a neurotrophin receptor tyrosine kinase which is a potent regulator of hippocampal LTP (Minichiello, 2009).

Microglia play a crucial role in clearing debris from the stroke microenvironment through phagocytosis to enhance neuroprotection (Wang, Leak and Cao, 2022). This was evident as expression of FC-γ receptor-mediated phagocytosis pathway genes such as FCGR1, FCGR3, FCGR4, SYK, and PIP5K1C was found to be upregulated in microglia of stroke brain in the current study. Interestingly, expression of these genes was further upregulated upon HDACi treatment indicating that HDACi enhances microglial response to ischemia and capacity for clearing cellular debris, limiting secondary neurodegeneration and inflammation (Qiao, Liu and Qie, 2023). Overall, HDACi-mediated phagocytic activity of microglia potentially ameliorates exacerbated neuroinflammation in the penumbra of the stroke brain (Lai, Zhang and Wang, 2014; Belov Kirdajova *et al*., 2020). Spatially, these transcripts in hippocampal microglia emphasize their role in mitigating damage and promoting recovery in regions critical for cognitive functions.

The microglial transcriptomic changes induced by HDACi in different penumbral regions of the stroke brain showcase a shift in microglial functionality from a proinflammatory phenotype to a phenotype which improves brain repair and regeneration by enhancing phagocytosis and reducing neuroinflammation.

This study also shows that HDACi altered the neuronal spatiotranscriptome to promote resilience and plasticity within the ischemic brain environment (Ganai, Ramadoss and Mahadevan, 2016; Kitahara *et al*., 2021). The inverse expression patterns in neurotrophic and neuroprotective pathways of these neurons in sham and stroke brains suggest broader implications for brain plasticity through upstream epigenetic mechanisms aimed at counterbalancing ischemia-associated excitotoxicity and neuroinflammatory response to promote tissue repair and reduce neuronal susceptibility to reperfusion injury (Delcuve, Khan and Davie, 2012; Kitahara *et al*., 2021).

In conclusion, our high-throughput spatio-transcriptomic analysis unravels the mechanisms by which a) microglia contribute to ischemic pathology with region-specificity and b) HDACi ameliorates reperfusion mediated ischemic-injury to neurons through modulation of microglial transcriptome. These findings emphasize an interplay between epigenetic regulation and spatial transcriptomic changes altering communication networks between microglia and neurons. Understanding the molecular mechanisms by which HDACi induced phenotypic changes together with altered spatial transcriptomic landscape in microglia of stroke brain would provide insight into development of potential microglia-mediated therapeutic strategies for brain recovery post ischemia.

## Acknowledgements

This article has been published on a preprint server: bioRxiv 2024.08.08.607139; doi: https://doi.org/10.1101/2024.08.08.607139 Components of Figure 1 have been created with BioRender.com.

## Author Contribution

Data acquisition: **KJ, RK**. Data analysis and interpretation: **KJ, ST.** Study design and conceptualization: **ST.** Writing of the original draft: **KJ, ST.** Reviewing & editing of manuscript: **all authors.** Supervision: **ST.**

## Data Availability

High-throughput, next-generation sequencing data from spatial transcriptomic libraries in this study is available at NCBI Sequence Read Archive (SRA) under BioProject PRJNA1144233.

## Ethics Approval

All experimental procedures performed *in vivo* were approved by the National University of Singapore Animal Care and Use Committee, under protocol NUS/IACUC/R19-0596.

---

**Figure 1 Supplementary Figure 1**

TTC (2,3,5-Triphenyltetrazolium Chloride) treated mouse brain sections across biological groups showcasing extent of infarct through viable (stained, red) tissue and non-viable (unstained, white) tissue. (n=3 in each experimental group).

**Figure 1 Supplementary Video 1**

Assessment of ischemic stroke affect through preliminary neurological deficit scoring (pNDS); video capture of each biological group showcases mild-moderate immobility and circling behaviour in the MCAo group while sham and HDACi-treated groups exhibit reduced to nil locomotor defects such as circling and difficulty reaching enrichment shell within home cage.

**Figure 2 Supplementary Video 1**

Video capture of IMARIS 9.5 reconstructions (30µm thick, z-stacked sections), showcasing surface creation and segmentation of labelled microglial cells (IBA1, AF488, false-colored red) that are processed to obtain morphometric data such as surface area and cellular volume to evaluate phenotypic status of microglia in the stroke brain after HDACi treatment (n=3 in each experimental group)

**Figure 3 Supplementary Figure 1**

(A,B,C) Tilescan (10x) images of complete brain sections across biological groups; (D,E,F) showcasing hippocampal expression of H3K27me3 (AF647, false-colored green) in microglia stained with IBA1 (AF488, false-colored red) that is upregulated upon ischemic stroke and ameliorated upon HDACi-treatment.

**Figure 3 Supplementary Video 1**

Video capture of (30µm thick, z-stacked sections) microglia stained with IBA1 (AF488, false-colored red) and H3K27me3 (AF647, false-colored green) across all three biological groups, showcasing elevation of histone mark expression and presence within the nuclei of microglial cells.

**Figure 4 Supplementary Figure 1**

(A,B) Tissue optimization timepoint experiments for brain samples with 10x Genomics Visium-TO slides for evaluation of optimum cDNA permeabilization time points (10x UG CG000238-RevE) prior to proceeding with gene expression and library preparation of Visium fresh-frozen spatial libraries. 6 minutes of permeabilization was selected based on optimization time points evaluated. (C) Antibody titre evaluation for IF-staining of spatial tissue sections (10x UG CG000312-RevD) identified concentration of antibodies (table in methodology).

**Figure 4 Supplementary Figure 2**

Permeabilized Visium fresh-frozen slides and quality control (QC) of RNA integrity checks and pre-cDNA amplification/ cleanup for visium libraries prior to preparation and sequencing.

**Figure 5 Supplementary Figure 1**

Alternative enriched (KEGG) pathways from commonly dysregulated genes of microglial spatiotranscriptomes across all 4 ROIs (hippocampus, thalamus, striatum, cortex). These pathways present additional targets and mechanisms that may be dysregulated throughout penumbral microglia after ischemic stroke and HDACi-administration.

**Figure 6 Supplementary Figure 1A**

Pathways enriched in HDACi-rescued, region-specific microglial spatiotranscriptomes (hippocampus-A, thalamus-B, striatum-C, cortex-D). Evaluation of pathways rescued by HDACi after ischemic injury, highlights pathways crucial for neuroprotective, neuroinflammatory and phagocytosis mediation responses.

**Figure 6 Supplementary Figure 1B**

Gene Ontology analyses of region-specific, microglial spatiotranscriptomes (hippocampus-A, thalamus-B, striatum-C, cortex-D) exhibit differential GO terms in respective compartments (BP, CC and MF) involved in HDACi-mediated rescue after ischemic injury in microglia.

**Figure 6 Supplementary Figure 2**

Heatmaps representing all dysregulated genes within selected biological pathways (HDACi vs MCAo Log2fc difference>1.0, Euclidean distance metric with hierarchical clustering through complete linkage) in each penumbra microglial spatiotranscriptomes (hippocampus-A, thalamus-B, striatum-C, cortex-D). Microglial heterogeneity in expression patterns of these pathways further highlights the specificity of microglial responses to ischemic stroke and HDACi therapy based on spatial contexts within the mouse brain.

**Figure 6 Supplementary Figure 3**

Circular Network Plot (cnetplot) of HDACi-rescued pathways and targets in each microglial spatiotranscriptome (hippocampus-A, thalamus-B, striatum-C, cortex-D) post-ischemic stroke. Genes (nodes) are positioned around the plot, sized by log2 fold change (log2FC) and colored from red (upregulated) to blue (downregulated). Pathway nodes are centrally located, sized by the number of associated genes, and color-coded by functional categories. Key pathways identified in the study including “MAPK signaling,” “PI3K-Akt signaling,” and “Fc gamma R-mediated phagocytosis.” are depicted with lines indicating gene-pathway associations, illustrating the molecular interactions affected by HDACi treatment.

**Figure 2. Supplementary File 1**

Morphometry data table consisting of IMARIS *Surfaces* feature metrics (Volume, Surface Area of microglial cells)

**Figure 6. Supplementary File 1**

Enriched Kyoto Encyclopedia of Genes and Genomes (KEGG) pathways of interest from individual microglial spatiotranscriptomes.

